# Dynamical localization of a thylakoid membrane binding protein is required for acquisition of photosynthetic competency

**DOI:** 10.1101/199943

**Authors:** Andrian Gutu, Frederick Chang, Erin K. O‘Shea

## Abstract

Vipp1 is highly conserved and essential for photosynthesis, but its function is unclear as it does not participate directly in light-dependent reactions. We analyzed Vipp1 localization in live cyanobacterial cells and show that Vipp1 is highly dynamic, continuously exchanging between a diffuse fraction that is uniformly distributed throughout the cell and a punctate fraction that is concentrated at high curvature regions of the thylakoid located at the cell periphery. Experimentally perturbing the spatial distribution of Vipp1 by relocalizing it to the nucleoid causes a severe growth defect during the transition from non-photosynthetic (dark) to photosynthetic (light) growth. However, the same perturbation of Vipp1 in dark alone or light alone growth conditions causes no growth or thylakoid morphology defects. We propose that the punctuated dynamics of Vipp1 at the cell periphery in regions of high thylakoid curvature enable acquisition of photosynthetic competency, perhaps by facilitating biogenesis of photosynthetic complexes involved in light-dependent reactions of photosynthesis.

## INTRODUCTION

Oxygenic photosynthesis is a metabolic process that must be dynamically modulated to accommodate varying light conditions and cellular needs (Eberhard *et al.*, 2008). If extraneous light energy is not productively engaged in a timely manner, phototoxic damage can arise and lead to cellular death (Apel and Hirt, 2004). These light-dependent reactions are catalyzed by photosynthetic protein complexes that are assembled in a specialized membrane system which forms the thylakoid compartment in both chloroplasts and cyanobacterial cells (Rast *et al.*, 2015). Although much is known about the structure and function of individual photosynthetic complexes, it is still unclear how the thylakoid membrane system is formed or maintained, and how the photosynthetic complexes are assembled in the thylakoid membrane (Pribil *et al.*, 2014; Rast *et al.*, 2015). These two issues are interrelated – thylakoid structure depends on the presence of photosynthetic complexes (Pribil *et al.*, 2014; Zhang *et al.*, 2014), and assembly of photosynthetic complexes depends on the presence of thylakoid membrane (Nickelsen and Rengstl, 2013; Yang *et al.*, 2015). Thus, experimentally uncoupling the factors required for thylakoid membrane formation from those required for photosynthesis is difficult.

Vipp1 (<u>V</u>esicle-<u>i</u>nducing <u>p</u>rotein in <u>p</u>lastids <u>1</u>), a conserved protein in plants, algae and cyanobacteria, has been proposed to play a role in thylakoid formation (Kroll *et al.*, 2001; Vothknecht *et al.*, 2012; Heidrich *et al.*, 2017). Vipp1 is not known to participate in photosynthetic reactions and its deletion in plants causes loss of photosynthesis and thylakoid organization, as well as loss of viability (Kroll *et al.*, 2001; Westphal *et al.*, 2001). The biophysical properties of Vipp1 are consistent with a membrane-related function – in vitro studies of Vipp1 revealed that it is a soluble protein with high affinity for lipid components (Otters *et al.*, 2013; McDonald *et al.*, 2015). On membrane surfaces prepared in vitro, recombinant Vipp1 forms higher order oligomers and, under certain conditions, mediates vesicle fusion (Hennig *et al.*, 2015). Thus, Vipp1 may facilitate thylakoid membrane formation, with one recent model proposing that oligomeric Vipp1 is an intermembrane lipid transporter, possibly transporting lipids from the plasma (or chloroplast inner) membrane to the thylakoid membrane (Heidrich *et al.*, 2017).

However, not all experiments are consistent with the above hypothesis. When expression of Vipp1 is reduced to a low level in cyanobacteria (Gao and Xu, 2009), photosynthetic output was severely diminished but thylakoid morphology remained intact, suggesting that Vipp1 plays a role in photosynthesis in addition to, or instead of, being required for thylakoid formation. This observation, combined with the fact that deletion of a core photosynthetic complex induces a thylakoid morphology defect (Zhang *et al.*, 2014), raises the possibility that Vipp1 may not directly facilitate thylakoid formation but instead its function may be related to photosynthesis. If this is true, knockdown or deletions of Vipp1 could generate a thylakoid morphology defect as a secondary consequence. Given the pleiotropy of *vipp1* phenotypes and the contrasting conclusions reached by other studies (Aseeva *et al.*, 2007; Fuhrmann, Gathmann, *et al.*, 2009), the role of Vipp1 in thylakoid membrane formation and in photosynthetic complexes formation remains unclear.

Previous microscopy work demonstrated that Vipp1 manifests as both diffuse and rare concentrated signals (Nordhues *et al.*, 2012; Zhang *et al.*, 2012; Bryan *et al.*, 2014). In cyanobacteria exposed to damaging high-light induced stress, the diffuse form of Vipp1 concentrates into long-lived immobile foci at the cell periphery, which were suggested to be important for light-induced stress protection (Bryan *et al.*, 2014). However, the native localization and dynamics of Vipp1 in normal (non-stressful) growth conditions has not been well characterized. Furthermore, without a perturbation that targets Vipp1 localization, the functional importance of this localization for photosynthesis or thylakoid membrane formation is not clear.

Here we show that in normal growth conditions, fluorescently labeled Vipp1 is highly dynamic, continuously exchanging between two fractions – a punctate fraction at the cell periphery that is concentrated at high curvature regions of the thylakoid, and a diffuse fraction that is uniformly distributed in the cytoplasm. By rapidly perturbing the spatial distribution of Vipp1 in living cells, we show that native Vipp1 localization is not required for thylakoid membrane formation, but is essential during the transition from non-photosynthetic to photosynthetic metabolism. We propose that the punctate fraction of Vipp1 is the cytological manifestation of the oligomeric and membrane-bound forms of Vipp1, whose role is to enable acquisition of photosynthetic competence. We hypothesize that Vipp1 facilitates the functional assembly of photosynthetic complexes.

## RESULTS

### Vipp1 forms transient puncta at regions of high thylakoid curvature at the cell periphery

To gain insight into the cellular role of Vipp1 we studied its localization and dynamics in live cells of *Synechocystis* sp. PCC 6803. To fluorescently label endogenous Vipp1 we integrated a Vipp1-mGFPmut3 fusion construct (hereafter Vipp1-GFP) at the native *vipp1* locus via homologous recombination. Both the expression level of Vipp1-GFP protein and the bulk growth rate of the strain harboring the construct were similar to that of the wild type parental strain (Fig. S1A-B), as was reported previously (Bryan *et al.*, 2014). In actively growing cells (doubling time ∼7 h) maintained at moderate light intensity (Fig. S1B), Vipp1-GFP is present in two fractions: as fluorescent diffraction-limited spots (hereafter puncta), and as a diffuse signal in the cytoplasm (Fig. 1A and Fig. S1C). Since Vipp1 puncta form when Vipp1 is fused to a known monomeric GFP variant that is least prone to induce multimerization of various bacterial targets (Landgraf *et al.*, 2012) and also when fused to the SNAP-tag (Keppler *et al.*, 2003) (Fig. S1D), which is not known to induce artificial clustering, we conclude that the observed Vipp1 puncta formation and diffuse signal reflect native spatial localization.

**Fig. 1.**
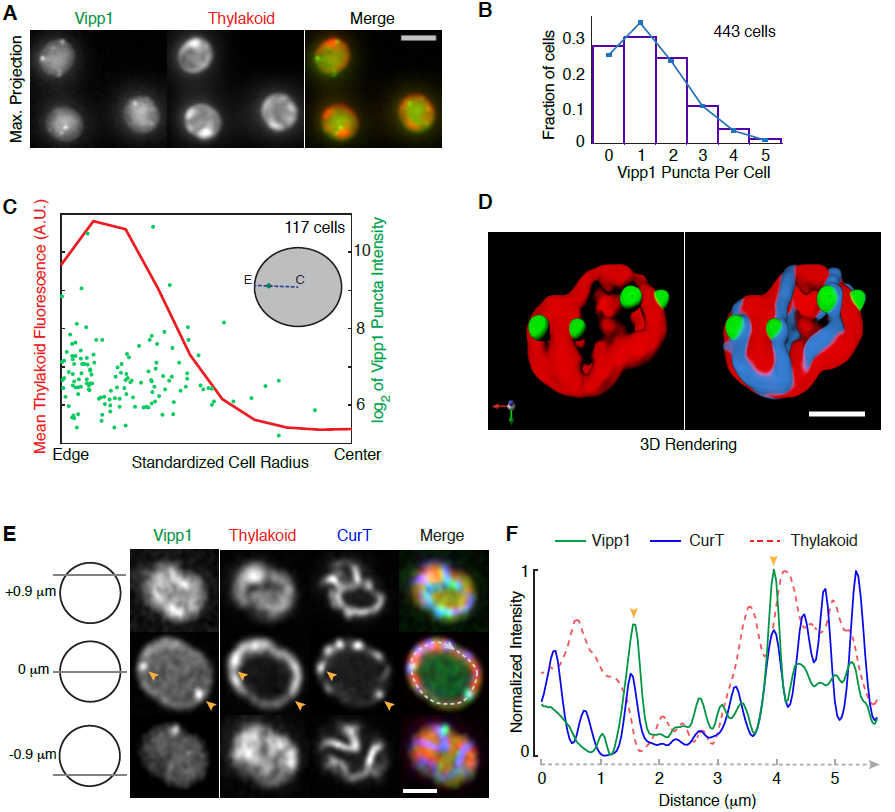
Vipp1 forms peripheral puncta that are located at regions of high thylakoid curvature. **A**) Representative live-cell epifluorescence images in the Vipp1 (GFP) and the thylakoid (far-red) channels. Images represent Maximum Z-Intensity Projections of 3D Z-stacks. Scale bar = 2 µm. **B**) Distribution of the number Vipp1 puncta per cell in a population of cells obtained from an exponentially growing culture. The mean and the variance of the distribution is 1.36. Puncta were identified by thresholding and segmenting filtered 3D Z-stacks. **C**) Analysis of the positioning of Vipp1 puncta relative to the cell radius. For each Vipp1 puncta identified at a given mid-cell plane (see inset diagram for an illustration), the corresponding distances to the cell center (labeled as “C “) and to the closest cell edge (labeled as “E “), as defined by brightfield segmentation masks, were obtained. The sum of the two distances defined the radius (blue dashed line) that was scaled from 0 to 1 on the x-axis. On the left y-axis, the mean of the thylakoid signal intensity along the line profiles (i.e. radii) connecting the cell centers, the Vipp1 puncta and the closest cell edge points is shown. The right y-axis shows the range of intensities (sizes) of all Vipp1 puncta analyzed. To limit the variability in size coming from dividing cells only cells whose area shape met an eccentricity value of 0.6 or less were analyzed. **D**) 3D rendering of a super-resolution image of a live cell of *Synechocystis* PCC6803 expressing Vipp1-GFP (green) and CurT-mTurquoise2 (blue). Thylakoids (red) are distributed at the cell periphery which by fluorescence microscopy show up as peripheral sheets. Vipp1 puncta are localized at the edge of thylakoid enrichments, at the same regions where the thylakoid membrane protein CurT is concentrated. Images were obtained with a laser scanning confocal system equipped with an Airyscan detector (Zeiss LSM880) which affords increased spatial resolution. Bar = 1 µm. For a full rendering of the same cell see Video S1. **E**) Representative live-cell confocal fluorescence image of a cell expressing both Vipp1-GFP and CurT-CFP obtained by Airyscan imaging. As diagramed on the left, the rows show Vipp1, thylakoid and CurT channels at the top, middle and the bottom of a cell. Two Vipp1 puncta are shown at mid-cell slice (orange arrowheads) which co-localize with CurT enrichments and with gaps in the thylakoid signal. A profile line (dashed in light grey) running circumferentially through the peripheral thylakoids was used to extract the intensities of Vipp1, CurT and thylakoid fluorescence and plotted in panel F. Bar = 1 µm. **F**) Intensities of the Vipp1, CurT and thylakoid signals along the mid-cell circumferential curved line profile traced in panel E. Raw signal intensities were normalized from 0 (minimum) to 1 (maximum) on the y-axis. Orange arrowheads indicate colocalization of Vipp1 puncta with CurT, which is enriched at regions of high thylakoid signal changes (edges).

In live cells growing in light on the microscope stage (see Methods), we find that the number of Vipp1 puncta per cell is well-described by a Poisson distribution with a mean and variance of 1.36 (Fig. 1B), suggesting that the formation of each punctum is an independent event. By measuring the positioning of each punctum relative to the cell boundary we find that most Vipp1 puncta localize near the cell periphery (Fig. 1C and Fig. S1E). At this region, the thylakoids are highly abundant (Fig. 1C), as estimated by the fluorescence emitted by the endogenous photosynthetic proteins in the far-red portion of the visible spectrum (Vermaas *et al.*, 2008). Moreover, within the thylakoids, we observed that Vipp1 puncta tended to localize to regions where the thylakoid signal is low (Fig. 1A and Fig. S1C). These low thylakoid signal regions correspond to the edges of the thylakoid stacks and are known as zones of high thylakoid membrane curvature (Heinz *et al.*, 2016). To confirm localization at sites of high thylakoid curvature, we asked whether Vipp1 puncta co-localize with CurT, a membrane protein enriched at these regions (Heinz *et al.*, 2016). For this, we used super-resolution imaging to determine the localization of Vipp1, CurT, and the thylakoid signal in living cells (see Methods). We find that in any given cell, Vipp1 puncta localize at the edge of thylakoid stacks where CurT is concentrated (Fig. 1D-F, Video S1). To quantify the relationship between Vipp1 puncta and CurT, we extracted and compared the intensity profiles of 76 arc lines that are each centered on a Vipp1 punctum found at the mid cell plane (Fig. S2A-B). We found that in general, Vipp1 signal positively correlated with the CurT signal and negatively with the thylakoid signal (Fig. S2A-B). We also used automatic 3D object-based colocalization analysis to evaluate the spatial relationship between Vipp1 puncta with CurT enrichments and find that the majority of puncta overlap with the volume of CurT objects (Fig. S2C-D). Based on these observations we conclude that in live cells the punctate fraction of Vipp1 localizes at cell periphery at or near regions of high thylakoid membrane curvature.

To better localize Vipp1 at the cell periphery and determine its relation to the thylakoid membranes we used immunoelectron microscopy. For immunodetection we used anti-GFP primary antibodies and gold-conjugated secondary antibodies to stain ultrasections (60 nm thickness) of freeze-substituted cells obtained from *vipp1-gfp* and wild type strains. We find that immunogold signals specific for Vipp1-GFP were enriched near the edges of thylakoids (high curvature regions which appear as tips in 2D representations) which typically converge near the plasma membrane (Fig. 2 and Fig. S3). Even though the antibodies can only bind the antigens from the surface of ultrasections, and therefore probe only a sparse subset of Vipp1-GFP, occasional clusters of 2-3 nanogold signals were observed near the thylakoid edges (Fig. 2C, Fig. S3A, Fig. S3F). This distribution is consistent with the idea that a fluorescent Vipp1 punctum consists of multiple Vipp1-GFP molecules closely juxtaposed in space. Combining the fluorescent and electron microscopy observations we conclude that the regions of high thylakoid membrane curvature are the sites of Vipp1 puncta localization.

**Fig. 2.**
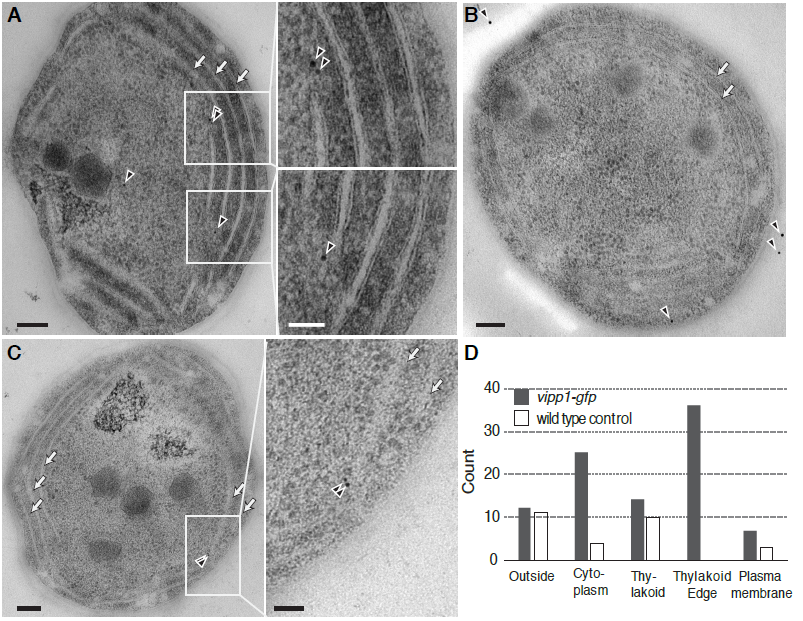
Detection of Vipp1-GFP by immuno-electron microscopy. Representative immunoelectron microscopy image of *vipp1-gfp* (**A** and **C**) or wild type cells (**B**) stained with anti-GFP primary antibodies detected by gold-conjugated secondary antibodies (black arrowheads). Arrows show the stacked thylakoids which run parallel to the plasma membrane. Scale bar = 100 nm (inset scale bar = 50 nm). **C**) Count of gold particles categorized by the proximity to the nearest cellular feature (plasma membrane, thylakoid edges, thylakoid (excluding edges), cell outside and the cytoplasm) in 35 independent whole cell sections obtained from either *vipp1-gfp* or wild type cells, respectively.

At a given time, many cells appear to contain no Vipp1 puncta (Fig. 1B), raising the question of whether only a fraction of cells are capable of forming Vipp1 puncta or if all cells can form puncta that are relatively short lived. We used time-lapse fluorescent microscopy to investigate Vipp1 dynamics in cells growing photosynthetically on the microscope stage and find that sparse Vipp1 puncta dynamically appear and disappear in all cells over time (Video S2). Furthermore, we also observed that in a small minority of cells (under 5%), Vipp1 is present exclusively as very bright and stable puncta, but these are observed only in non-growing cells (Fig. S4A). Thus, transient Vipp1 puncta likely exist in all growing cells over time, explaining the zero puncta bin seen in the distribution of Vipp1 counts per cell (Fig. 1B).

To determine if Vipp1 puncta form directly on the membrane or if they form in the cytoplasm and diffuse to the membrane, we imaged photosynthetically growing cells at high temporal resolution to capture the appearance, disappearance, and the movement of Vipp1 puncta. We find that the majority of Vipp1 puncta rise and fall in intensity on a time scale of 1-2 min (Fig. 3A-B, Fig. S4B, Video S3) and their mobility at the cell periphery is greatly limited (Fig. 3C and Video S4). This constrained mobility and the rise and fall in fluorescence within a diffraction limited volume is consistent with a Vipp1 assembly process occurring on the membrane. Given that Vipp1 binds membranes as a homo-oligomeric complex in vitro and exists as a distribution of oligomeric states in cell lysates (Fuhrmann, Bultema, *et al.*, 2009; Hennig *et al.*, 2015; McDonald *et al.*, 2015; Heidrich *et al.*, 2016), we infer that the dynamically forming Vipp1 puncta, occurring at or near the highly curved regions of the thylakoid compartment, represent events of punctuated oligomerization and de-oligomerization between its cytosolic and its membrane-bound forms.

**Fig. 3.**
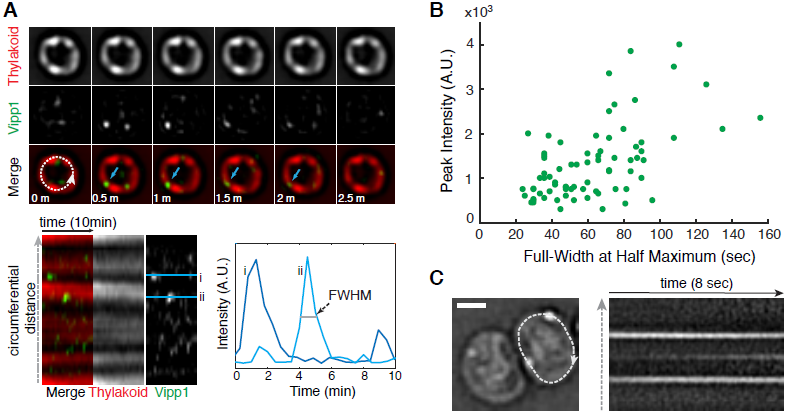
Vipp1 forms transient puncta of limited mobility at the cell periphery. **A**) Time-series montage of a live cell growing photosynthetically on the microscope stage. The images were acquired every 30 seconds in the Vipp1 and thylakoid channels for a duration of 10 minutes. The montage shows only the mid-cell Z plane for the first 2.5 minutes. The blue arrows highlight the appearance and disappearance of a Vipp1 puncta in consecutive time points. The dashed circumferential line was used to extract the intensity profiles of the two channels and build the kymogram displayed below the montage. The Vipp1 puncta appear and disappear at the edge the thylakoids which manifest as horizontal streaks. Two examples of Vipp1 intensity profiles in time (i and ii) are graphed on the right to illustrate how the full-width at half-maximum (FWHM) durations and peak intensities were obtained for a given punctum. **B**) Distribution of Full-Width at Half Maximum (FWHM) durations relative to their peak intensities for a representative subset of Vipp1 puncta. The values were extracted from the individual kymograms of 15 growing cells that were imaged every 30 seconds for a duration of 20 min (see examples in Video S3). **C**) Example of Vipp1 puncta mobility recorded by continuous imaging (frame rate 81 milliseconds). On the left, the Laplacian-of-Gaussian filtered image of a cell containing two peripheral Vipp1 puncta is shown. A circumferential line profile traversing the cell periphery (dashed line) was used to extract the kymogram on the right to illustrate the constrained mobility of Vipp1 puncta over time. The kymogram also captures the birth of a new Vipp1 punctum as evidenced by the appearance of a middle horizontal streak over time. See Video S4 for examples of other cells imaged at similar time scale resolution. Scale bar =1 µm.

### Spatio-temporal distribution of Vipp1 enables photosynthetic competency

To test the functional importance of the spatial distribution and dynamics of Vipp1 in the cell, we designed a perturbation in which Vipp1 was rapidly and reversibly localized to the cell center, away from the cell periphery, without changing its overall concentration. To achieve this perturbation we used the anchor-away technique (Liberles *et al.*, 1997; Haruki *et al.*, 2008) in which a protein of interest can be rapidly moved away from its native location by inducing a drug-dependent dimerization to another protein, known as the anchor, that is in a different sub-cellular location (Fig. 4A). For the anchor we used the histone-like HU protein that binds to DNA in the nucleoid and is localized at the center of the cell (Wang *et al.*, 2011). In the presence of the drug rapamycin, cells expressing the components of the anchor-away system rapidly alter the native localization of Vipp1 by coalescing the entire pool of Vipp1 to the nucleoid region of the cell, away from the cell periphery (Fig. 4B, Fig. S5). We note that this perturbation of Vipp1 localization inherently alters the dynamics of puncta formation occurring at the cell periphery.

**Fig. 4.**
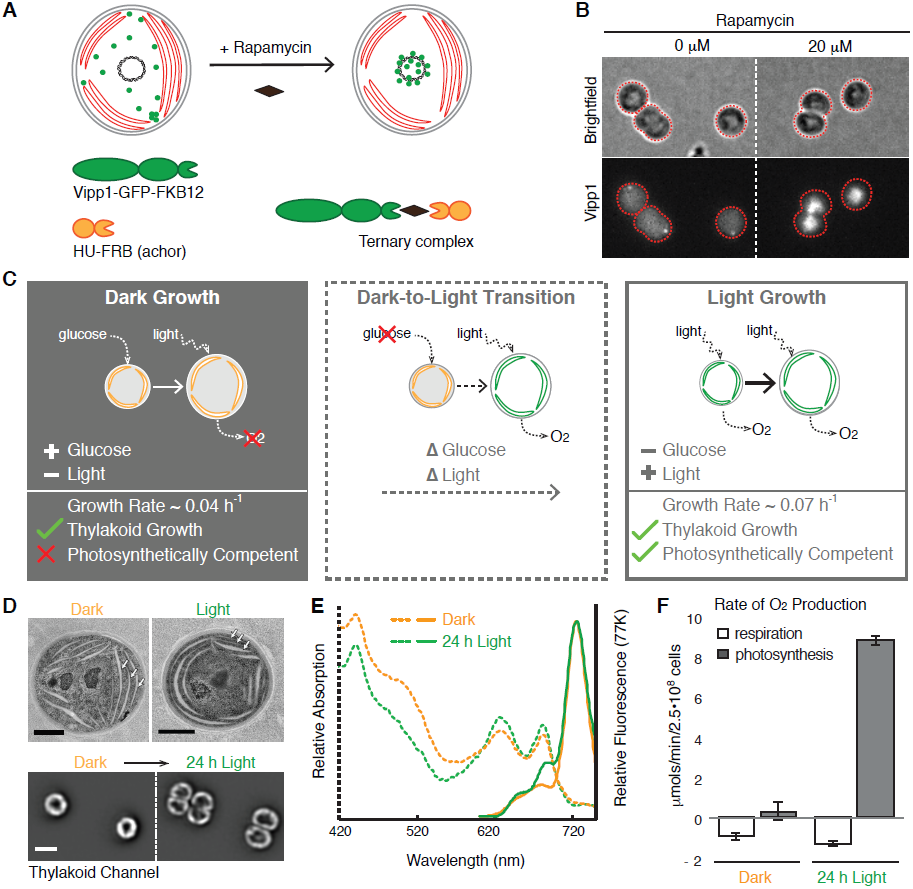
Perturbation of Vipp1 localization and characterization of the growth conditions in which the perturbation was applied. **A**) Diagram illustrating how Vipp1 can be relocalized to the nucleoid, away from the cell periphery using the anchor-away approach. A schematic cell is shown, with red disks representing thylakoids, the black circle in the center representing the nucleoid, and green dots representing the diffuse and punctate fractions of Vipp1. Human FKBP12 (12 kDa FK506-binding protein) was fused to Vipp1-GFP and the FRB (FKBP-rapamycin binding) domain was fused to the histone-like HU protein. In the presence of the cell permeable drug rapamycin, the Vipp1-GFP-FKBP12 chimera tethers to HU-FRB and forms a ternary complex enriched at the nucleoid due to the high binding affinity of HU to DNA. **B**) Rapamycin-dependent relocalization of Vipp1 to the nucleoid region at the cell center is effective and rapid. Raw images of Maximum Z-Intensity Projections in the brightfield and Vipp1 channels of exponentially growing cells expressing the Vipp1-GFP anchor-away system in the presence or absence (solvent only) of rapamycin after 20 min incubation. Nucleoid-localized Vipp1-GFP manifests as bright concentrations Vipp1-GFP signal at the cell centers. The grayscale contrast is the same for both conditions. Cell outlines based on the brightfield channel are overlaid in red. **C**) The three growth conditions used for testing the functional importance of native Vipp1 localization via anchor-away: *Dark Growth*, based on glucose, during which thylakoids are multiplied while photosynthetic competency is highly reduced; *Dark-to-Light Transition*, during which dark-grown cells are shifted to light in glucose-free media so that cells can regain their photosynthetic competency over time (“delta “ symbols in front represent changes: glucose drop out and); and *Light Growth*, during which the cells grow in constant photosynthetic conditions. In each panel, a cell is shown growing from a smaller size to a larger size, and thylakoids are shown in orange if not competent for photosynthesis and in green if they do support photosynthesis. Photosynthetic competency is illustrated as the ability to produce O_2_ when light is applied (dotted arrows). Approximate growth rates shown were estimated by monitoring the optical density of liquid cultures and are consistent with previously published measurements (Anderson and McIntosh, 1991). **D**) Thylakoids are present in both dark-and light-grown cells. **Top** - representative electron microscopy images of dark-and light grown cells showing a similar thylakoids arrangement consisting of several cell peripheral stacks of 2-4 thylakoid sheets (arrows). Scale bar = 0.5 µm. **Bottom** - thylakoid fluorescence images of two dark-grown cells before (left) and after 24 hours of growth in light (right) directly on the microscope stage in glucose-free media. In both conditions, the thylakoid signal is enriched at the cell periphery with occasional gaps in fluorescence intensity that correspond to the thylakoid stacks’ edges. The same grayscale contrast applied for both conditions. Only the mid-Z planes of filtered image stacks are shown. Scale bar = 2 µm. The dark-grown cells were obtained by growing a culture in darkness on glucose for 144 h (∼ 8 generations), as described in Methods. **E**) Both dark- and light-grown cells contain photosynthetic complexes and pigments. Whole-cell absorption (dashed lines) and low-temperature (77K) fluorescence emission (solid lines) spectra of dark-grown (144 h in darkness) and light-grown (24 h post-dark) cultures. The chlorophyll (445 and 680 nm) and the phycocyanin (625 nm) peaks in the absorption spectra are present in both growth conditions, albeit at slightly lower level in dark-grown culture. The fluorescence emission spectra (excitation at 445 nm) reveal the signature peak of the photosystem I at 725 nm present in both cultures, and the peaks associated with active photosystem II (685 nm and 690 nm) which appear conspicuous in the light-grown culture only, as was reported previously (Barthel *et al.*, 2013). **F**) Dark-grown cells are photosynthetically incompetent. Rates of whole-cell oxygen production of dark-and light-grown cultures. Light-grown cells were obtained by shifting the dark-grown cells to light for 24 h in glucose-free medium. Linear rates of oxygen production were obtained from cells maintained in darkness (respiration, i.e. oxygen consumption) or in saturating light (photosynthesis, i.e. oxygen evolution) for the duration of the measurement. Respiratory activity which originates from the protein complexes that also localize in thylakoid membranes is similar between the dark-and light-grown samples. Error bars are standard error of the mean obtained from three biological replicates.

To reveal the functional role of native Vipp1 localization, we chose to apply the anchor-away perturbation in three different physiological conditions that enable us to disentangle interactions between thylakoid formation and photosynthesis – *Dark Growth*, *Dark-to-Light Transition*, and *Light Growth* (Fig. 3C). During *Dark Growth* (Anderson and McIntosh, 1991), cells rely on glucose from the media to grow and divide in the absence of photosynthetic light, while still multiplying their thylakoid membranes and associated pigments. However, dark-grown cells are not photosynthetically competent (Barthel *et al.*, 2013), meaning they are incapable of immediate oxygen production upon application of light. In the *Dark-to-Light Transition,* photosynthetic competency is induced when dark-grown cells are shifted to light conditions for at least 6 - 8 hours (Barthel *et al.*, 2013). During *Light Growth,* cells rely on photosynthesis to grow and divide in the absence of glucose, and also continuously multiply their thylakoids and photosynthetic components. We confirmed in our own experiments that dark-grown cells, even after multiple cell divisions, still maintain and produce thylakoids and photosynthetic pigments as estimated by microscopy, and spectrophotometry respectively (Fig. 4D-E). We note that in electron microscopy images, the thylakoid sheets in dark-grown cells appeared less tightly stacked than in light-grown cells, however they still maintained a peripheral arrangement that was also observed in fluorescence images of live cells, which later grew and divided when shifted to *Light Growth* (Fig. 4D). We also confirmed that dark-grown cells are photosynthetically incompetent as evidenced by their inability to immediately evolve oxygen in the presence of saturating light (Fig. 4F). When these dark-grown cells are shifted to *Light Growth* in glucose free media, the photosynthetic competency is restored within 24 h (Fig. 4F) as shown previously (Barthel *et al.*, 2013).

We asked whether the rapid drug-induced relocalization of Vipp1 to the nucleoid in the three conditions (*Dark*, *Dark-to-Light* and *Light*) would elicit either a growth or thylakoid morphology defect. For each of the conditions, we added rapamycin to induce Vipp1 relocalization and used time-lapse fluorescent microscopy to monitor thylakoid content, Vipp1 localization and cell growth for at least half of the cell cycle (Fig. 5A-C). For longer timescales, we measured growth by monitoring turbidity in bulk culture (Fig. 6A-B) and examined thylakoid morphology by both fluorescence and electron microscopy (Fig. 6C). We find that Vipp1 relocalization causes no significant growth or thylakoid morphology defects in *Dark* or *Light* conditions (Fig. 5A-B, Fig. 6A-C, Fig. S6), but causes a severe growth defect when Vipp1 is relocalized during the *Dark-to-Light Transition* (Fig. 5C, Fig. 6A, Video S5). The growth defect observed during the *Dark-to-Light Transition* is not due to nucleoid-related changes or to drug addition, since a control strain in which heterologously-expressed GFP is relocalized to the nucleoid exhibits no growth phenotype (Fig. 6B). The severity of the growth defect correlates with the timing of Vipp1 relocalization during the *Dark-to-Light Transition,* with less of a growth defect observed when the perturbation occurs later in the transition (Fig. S7A). Intriguingly, the most severe growth defect is observed when relocalization is induced at the start of the *Dark-to-Light Transition,* which coincides with the peak burst in Vipp1 puncta formation in normal conditions (Fig. S7B, Video S6).

**Fig. 5.**
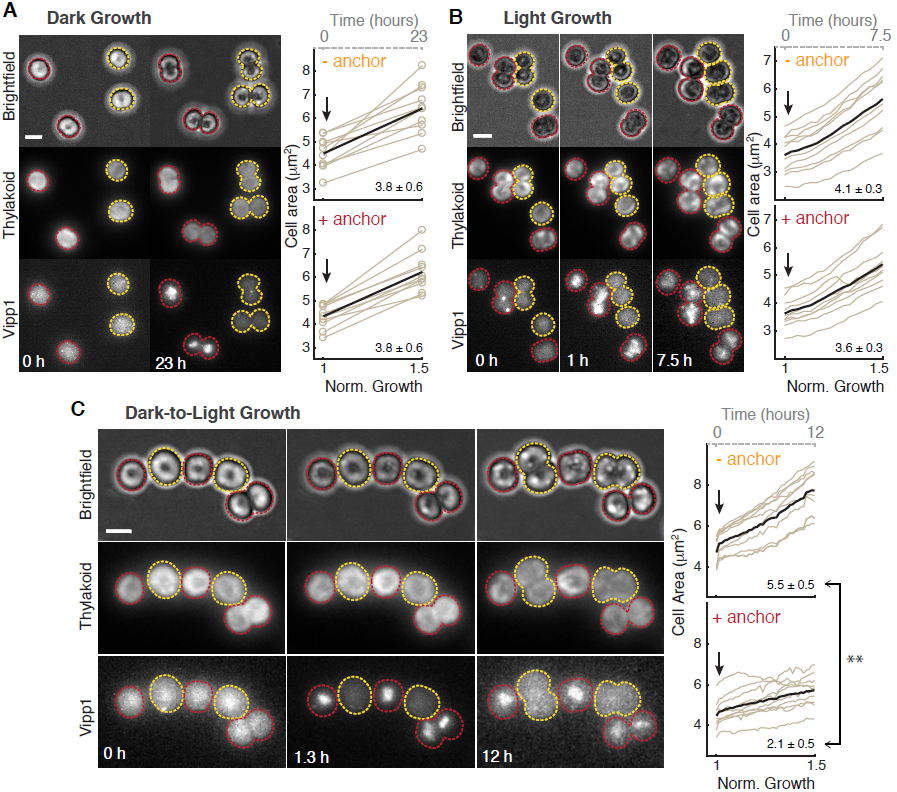
Relocalization of Vipp1 to the nucleoid has no significant effect on *Dark* or *Light Growth* but elicits a severe growth defect during *Dark-to-Light Transition*. **A**) Relocalization of Vipp1 induces no growth defect during *Dark Growth*. Left: Example images of dark-grown cells (Maximum Z-Intensity Projections, 12 Z-steps) in brightfield, thylakoid and Vipp1 channels before addition of rapamycin (0 h) and after 23 hours of growth in darkness. A mix of two strains are shown, one (yellow outlines) in which the Vipp1-GFP-FKBP12 construct is expressed alone, and another (red outlines) in which the HU-FRB anchor construct is also expressed *in trans* (i.e. responsive to rapamycin-based relocalization). 20 µM rapamycin was added to the agarose pad holding the cells on the microscope stage immediately after time 0 h. Right: quantification of cell growth as areas (extracted from automatic segmentations in the brightfield channel - open circles) over time of ten random cells obtained from each strain (with or without HU-FRB anchor) in the same field of view. On the bottom x-axis, normalized time is shown as the time it took for “-anchor “ cells to increase the average cell area by 50 %. On the top x-axis (dashed grey line), time is shown in hours. Arrows indicate the addition of rapamycin to the agarose pad. Average growth trace is shown as a black bold curve. The mean and standard error of the mean of area expansions rates of the 10 cells are shown in the lower right corners. We found no significant difference between the means of “+anchor “ and “- anchor “ cells as calculated with the two-sample two-tailed *t-test* at 5% significance level. **B**) Relocalization of Vipp1 induces no significant growth defect during *Light Growth*. Similar to panel A - on the left: selected images of light-growing cells obtained from a timelapse movie (8 Z-steps, every 30 min for 7.5 hours total). Right: quantifications of cell growth as areas over time of ten random cells obtained from each strain from the same field of view. No significant differences between the means of “+anchor “ and “- anchor “ cells rates (shown at the lower right corner) as calculated with the two sample two-tailed *t-test* at 5% significance level were found. **C**) Relocalization of Vipp1 during *Dark-to-Light Transition* induces a severe growth defect. Similar to panel A - on the left: selected images of dark-grown cells shifted to light growth obtained from a timelapse movie (10 Z-steps, every 20 min for 12 hours). Photosynthetic light and rapamycin were applied immediately after the first timepoint. On the right: quantification of cell growth as areas over time of ten random cells obtained from each strain from the same field of view. The difference between the means of “+anchor “ and “- anchor “ cells rates (shown in the lower right corner) was found to be significant (* * P < 0.001) based on the two-sample two-tailed *t-test* at 5% significance level.

**Fig. 6.**
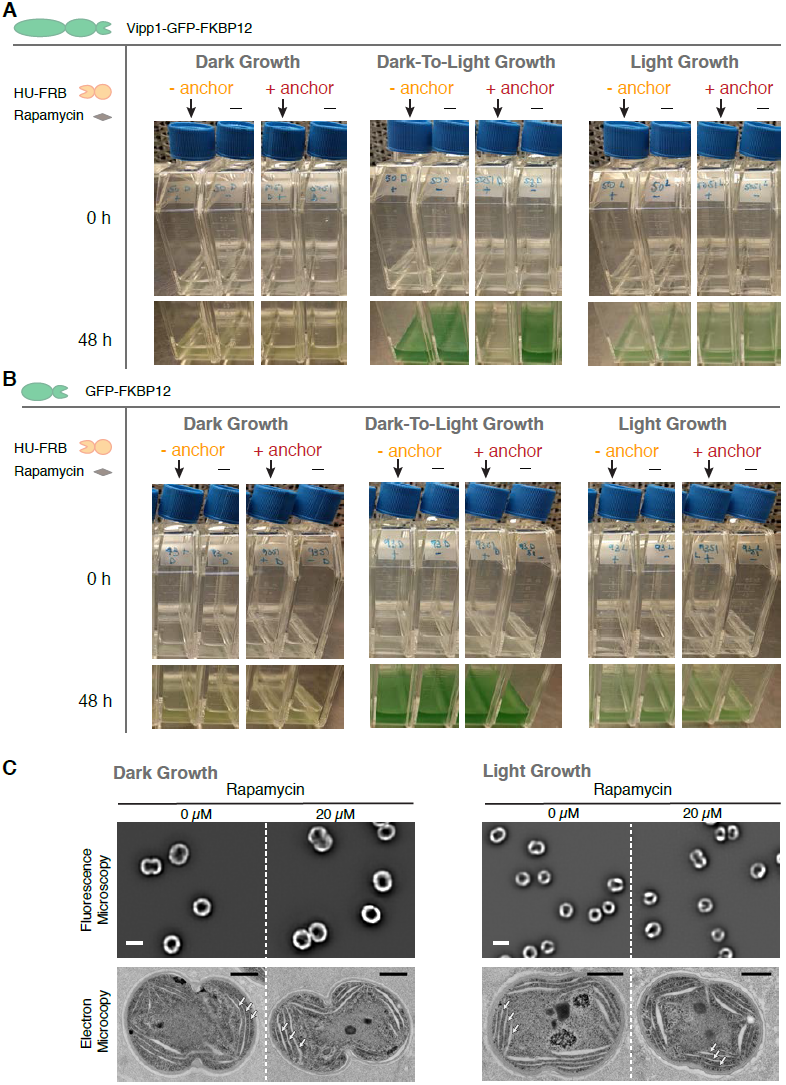
Vipp1 relocalization produces no growth or thylakoid morphology defect in *Dark* or *Light Growth* cultures but elicits a severe growth defect in the *Dark-To-Light Transition*. **A**) Effect of Vipp1 relocalization on growth in bulk culture. Dark- and light growing *vipp1-gfp-fkbp12* cells of the two strains - with or without the HU-FRB anchor construct expressed *in trans* - were resuspended in fresh media in pairs of flasks at the same starting inoculum and incubated in dark or in light for 48 h. 20 µM rapamycin (arrows) or solvent only (dimethyl sulfoxide, denoted as “- “) was added at time 0 h. **B**) Control cultures showing that relocalization of GFP alone at the nucleoid elicits no growth defects. Dark- and light growing *gfp-fkbp12* cells of the two strains - with or without the HU-FRB anchor construct expressed in trans - were inoculated in fresh media in pairs of flasks at the same starting inoculum and incubated in dark or in light for 48 h. 20 µM rapamycin (arrows) or solvent only (dimethyl sulfoxide, denoted as “- “) was added at time 0 h. **C**) Relocalization of Vipp1 during *Dark* or *Light Growth* induces no changes in overall thylakoid content or morphology. **Top** - filtered images in the thylakoid channel (mid-cell plane) obtained from cells expressing all the components of Vipp1 anchor-away in the presence or absence of rapamycin after 48 h of growth in darkness (left) or in light (right). The images are displayed at the same grayscale contrast for rapamycin and no rapamycin treated cells. The overall thylakoid morphology (i.e. peripherally localized signal with occasional gaps in fluorescence intensity that correspond to regions of high curvature of thylakoid membranes) in the rapamycin-treated cells is similar to that of the control cells. Scale bar = 2 µm. **Bottom** - corresponding electron microscopy images of representative cells from each of the conditions tested. Stacked thylakoids (arrows) are present in both dark- and light-grown cells and their morphology and arrangement remain unaffected by Vipp1 perturbation. Scale bar = 0.5 µm. See Fig. S6 for more electron microscopy examples of cells from these conditions.

Based on the following observations, we conclude that the native localization of Vipp1 is not necessary for thylakoid membrane growth, but is necessary for enabling acquisition of photosynthetic competency. First, since perturbing Vipp1 in the *Dark* condition (in which there is active thylakoid growth) triggers no defect in thylakoid growth or morphology (Fig. 6C, Fig. S6A-B), we conclude that Vipp1 localization is not required for thylakoid membrane growth. However, since perturbing Vipp1 in the *Dark-to-Light* condition (in which cells must enable photosynthesis in order to grow) elicits a severe growth defect (Fig. 5C, Fig. 6A), we infer that native Vipp1 localization is necessary to enable acquisition of photosynthetic competency. As photosynthetic competency is already established during *Light Growth* and Vipp1 perturbation does not trigger a growth or thylakoid morphology defect (Fig. 5B, Fig. 6C, Fig. S6C-D), we infer that Vipp1 localization is not critical for maintaining thylakoid membranes or supporting active and pre-existing photosynthetic capacity.

## DISCUSSION

The presence of Vipp1 as diffuse and punctate signals has been previously reported in cyanobacteria exposed to two extreme physiological conditions: low light (8 µE m^-2^ s^-1^ intensity) and high light (600 µE m^-2^ s^-1^ intensity) (Bryan *et al.*, 2014). In each of these conditions, both suboptimal for growth ((Kopecna *et al.*, 2012)), Vipp1 was predominantly distributed either as a diffuse signal in low light or as stable, long-lived puncta in high light (Bryan et al., 2014). Here we characterized in detail the localization and dynamics of these two Vipp1 fractions in exponentially growing cells, maintained at moderate light intensity (100 µE m^-^2 s^-^1 intensity), and show that Vipp1 continuously exchanges between the diffuse and punctate fractions on a timescale of minutes (Fig. 1C, Fig. 3A-B, Video S3). This punctate fraction is short-lived, displays limited mobility (Fig. 3) and localizes to regions of high thylakoid membrane curvature (Fig. 1D-F, Fig. S2). The localization of Vipp1 is consistent with the known biochemical properties of Vipp1 oligomerization and membrane binding (Fuhrmann, Bultema, *et al.*, 2009; Otters *et al.*, 2013; Hennig *et al.*, 2015), particularly the high affinity to membrane substrates manifesting high stored curvature elastic stress (McDonald *et al.*, 2015). Combining these observations with the spatiotemporal dynamics of Vipp1, we propose that transient formation of Vipp1 puncta at regions of high thylakoid curvature represent events of assembly and disassembly of Vipp1 oligomers that accompanies normal cellular growth.

The ability of *Synechocystis* PCC6803 cells to grow non-photosynthetically, while still maintaining and multiplying their thylakoids (Fig. 4D) – which are also the sites of respiratory activity (Mullineaux, 2014) – allowed us ask whether the localization and dynamics of Vipp1 is important for thylakoid membrane growth alone. By experimentally perturbing the native spatial distribution of Vipp1 in this *Dark* condition, we show that thylakoid membrane growth does not require native Vipp1 localization (Fig. 6C). This suggests that thylakoids (at least the ones generated in *Dark* growth) can be made even when Vipp1 is not available at the thylakoids or cell periphery, casting doubt on the hypothesis that Vipp1 is involved in lipid (or other components) transport from the plasma membrane to thylakoids (Heidrich *et al.*, 2017).

During the *Dark-To-Light Transition* the cells must either turn on pre-existing inactive photosynthetic protein components or synthesize new active ones. As native Vipp1 spatial distribution is critical in this condition (Fig. 5C, Fig. 6A, Video 5), and given the previously described requirement of Vipp1 to support photosynthesis (Gao and Xu, 2009b), we infer that in *Dark-to-Light* cells Vipp1 must be involved in the light-dependent synthesis or maturation of photosynthetic machinery. In other words, native Vipp1 localization enables the acquisition of photosynthetic competency when light is available. Since altering Vipp1 localization in *Light* condition elicits no significant effect on growth (Fig. 5B, Fig. 6A), we suspect that the Vipp1 perturbation is less effective because the photosynthetic competency is already established in these cells. Taken together, we conclude that native localization of Vipp1 is not required for thylakoid growth, but for enabling acquisition of photosynthetic competency.

Although relocalization of Vipp1 to the nucleoid alters both the diffuse and the punctate fractions, we postulate that the punctuated dynamics of Vipp1 (which are abolished during the relocalization experiment) at high curvature regions of the thylakoid are important for enabling the acquisition of photosynthetic competency. We base this on our observations that only in actively growing cells Vipp1 continuously exchanges between the diffuse and punctate fractions, in contrast to non-growing cells in which cellular Vipp1 remains “locked “ as stable puncta (Fig. S4). In addition, at the beginning of the *Dark-to-Light* condition when the need to initiate photosynthesis is high, the rate of Vipp1 puncta formation is also greatly increased as is the sensitivity of cells to Vipp1 relocalization (Fig. S7). Finally, Vipp1 puncta form at regions of thylakoids marked by CurT enrichments (Fig. 1D-F), a specialized zone previously proposed to act as biogenesis centers for new photosynthetic complexes (Heinz *et al.*, 2016). We note that the association between Vipp1 and CurT that we measured cytologically (Fig. S2) has not been confirmed in previous biochemical interaction studies (Bryan *et al.*, 2014; Heinz *et al.*, 2016), which suggests only an indirect connection between Vipp1 and CurT at these regions of the thylakoid.

What could be the molecular mechanisms by which Vipp1 enables acquisition of photosynthetic competency? Combining our analysis of Vipp1 localization and dynamics with its known biochemical properties, we hypothesize that punctuated oligomerization of Vipp1 at regions of high thylakoid curvature mediates functional light-dependent assembly of photosystem(s) complexes. Perhaps due to its membrane-binding and/or lipid rearrangement properties (Hennig *et al.*, 2015), the oligomerized form of Vipp1 could facilitate efficient translation and/or assembly of newly made photosynthetic complexes in the membrane – this may be accomplished by alleviating the membrane curvature elastic stress potentially emerging at thylakoid edges (McDonald *et al.*, 2015), assisting complex assembly by providing lipid cofactors, as previously proposed (Nordhues *et al.*, 2012; McDonald *et al.*, 2017), or mediating putative transient intermembrane contacts (Hennig *et al.*, 2015; Heidrich *et al.*, 2017). Our conclusions are consistent with the previous analysis of Vipp1 depletion in *Synechocystis*, which showed that when Vipp1 levels are lowered, the photosynthetic function is significantly reduced while respiration and thylakoid morphology remains unchanged (Gao and Xu, 2009). Similarly, deletion of Vipp1 in *Synechococcus* PCC 7002 implicated Vipp1 primarily in the translation or assembly of photosystem I, not for thylakoid membrane biogenesis per se (Zhang *et al.*, 2014). Additionally, Vipp1 has been shown to interact with specific chaperones, translation factors and core photosynthetic proteins, supporting a role for Vipp1 in the translation or assembly of photosystems (Bryan *et al.*, 2014).

Our results highlight the importance of Vipp1 dynamic behavior at high curvature areas of the thylakoid, which could act as sites of activation or synthesis of new photosynthetic complexes. Additionally, we present a new rapid perturbation of Vipp1 as a tool to biochemically probe which photosynthetic component(s) requires Vipp1 in its functional assembly. Finally, this work highlights how single-cell time lapse imaging and rapid perturbations can complement biochemical and genetic observations in understanding complex processes such photosynthesis that change in space and time.

## MATERIALS AND METHODS

### Strains and growth conditions

*Synechocystis* sp. PCC 6803 GT strain (kind gift of Dr. Wim Vermaas) (Trautmann *et al.*, 2012) was used for all the work presented. Strains were propagated on BG11-1.5 % GelRite (PlantMedia) containing 10 mM HEPES-KOH pH 8.0 in the presence of the appropriate antibiotic. Photosynthetically-grown cultures were grown at 30 °C either in Nalgene plastic flasks tilted at low angle on an orbital shaker (70 RPM) or in 40 ml glass tubes bubbled with 1% CO_2_-air mix. Philips cool fluorescent tube light bulbs (F40T12/841 Alto) provided the light whose intensity was set to 30 (when flasks were used) or 100 µE m^2^ s^-^1 (when air was bubbled) as measured with a LI-COR 190R sensor. Depending on the conditions, the doubling time in the log-phase ranged from 10 to 6 hours (see example of a typical growth curve in Fig. S1B). For all growth or imaging experiments, the cells were prepared and maintained in media with no antibiotics. Prior to imaging, cells were acclimated for ∼30 min on the microscope stage under a thin agarose pad made with BG-11 medium at 30 °C while illuminated by an external LED white light source (set to ∼100 µE m^2^ s^-^1 intensity light measured at the objective lens) (See Light Microscopy and Image Analysis Section below). To sustain non-photosynthetic growth (*Dark Growth*), the cultures were supplemented with 27 mM glucose, kept in darkness and stimulated with a daily 5 min exposure of 5 µE m^2^ s^-1^ light as was previously established to promote their viability (Anderson and McIntosh, 1991). For all experiments, the cultures were grown in dark up to 144 hours on an orbital shaker (70 RPM) and diluted to an OD_750_ of 0.1 when their OD_750_ reached 1 or above (doubling time ∼16 h). To limit any light effect during *Dark Growth* perturbation of Vipp1, the cultures were kept in continuous darkness for the duration of the experiment.

### Genetic manipulations

To obtain the GFP-tagged allele of Vipp1, we cloned into pBR322 between EcoRI and NcoI using Gibson Assembly (Gibson, 2011) a chimeric construct containing the upstream region and the open reading frame of *sll0617* translationally fused (SGGG linker) at the C-terminus to the codon-optimized variant of mGFPmut3 (Landgraf *et al.*, 2012), the putative native 3’ untranslated region of *sll0617* (68 bp), the spectinomycin resistance cassette (*aadA*) and a ∼ 1 kb homology regions downstream of *sll0617*. The resulting plasmid (pAGH42) was used to transform the naturally competent *Synechocystis* according to published procedures (Zang *et al.*, 2007). As for the *vipp1-gfp* construct, the SNAP-tagged version of Vipp1 - also codon optimized - was cloned and transformed into *Synechocystis* cells.

To make the Vipp1-GFP-FKBP12 construct for the anchor-away relocalization system (Liberles *et al.*, 1997) we fused the codon optimized fragment of the human FKBP12 to the C-terminus of GFP (GSGG linker) in the Vipp1-GFP construct described above using the Gibson assembly procedure. The resulting plasmid (pAGH50) was transformed into the appropriate *Synechocystis* strain to replace the native *vipp1* locus. No differences in growth or Vipp1 localization and dynamics were observed between the *vipp1-gfp-fkbp12* and *vipp1-gfp* strains.

To make the HU-anchor fusion, we designed a construct synthesized by Integrated DNA Technologies (Coralville, IA) in which the codon optimized FRB fragment- was fused to the C-terminus (GSG linker) of HU (encoded by *sll1712*). The DNA construct included the putative native promoter and 5’ untranslated region (265 nt) of the *sll1712* gene. At the 3’end, the 68 bp 3’ untranslated region of *vipp1* was used. The HU-anchor construct was cloned by Gibson assembly downstream of the kanamycin resistance cassette in a pBR322-based plasmid designed to recombine into the *slr0168* neutral site of the *Synechocystis* chromosome (Gao and Xu, 2009).

A GFP-FKBP12 control construct (pAGH93) was made similarly to *vipp1-gfp-fkbp12*, except that the L03 synthetic promoter was used to drive its expression (Huang and Lindblad, 2013).

CurT-mTurquoise2 construct was described previously (Heinz *et al.,* 2016).

In all cases, the colonies obtained after transformation were re-streaked onto fresh selection media and the genetic modification was confirmed by PCR amplification with primers flanking the locus of interest. For long-term storage, all *Synechocystis* strains were kept at −80°C in BG11 with 8% dimethyl sulfoxide. All vectors used in this study are [will be] deposited in the Addgene plasmid bank.

### Light microscopy and image analysis

All epifluorescence imaging was performed on a Zeiss AxioObserver.Z1 inverted microscope equipped with an environmental chamber, a Definite Focus module, a hardware-triggered Hamamatsu ORCA Flash4.0 V2+ camera and a Plan-Apochromat 63x/1.40 Oil DIC M27 (NA 1.4) objective. Unless noted, all images were acquired at Nyquist sampling rate as Z-stacks in Zen 2.1 software (Zeiss). Sola SE (Lumencor, Beaverton, OR) white light source was used to image Vipp1 and the thylakoid in the GFP and the far-red fluorescence channels (Zeiss Filter Sets 38HE and 50), respectively. All brightfield imaging was done in Köhler illumination with light passing through the same filter cube as the one used for Vipp1 channel.

For superresolution imaging we used Zeiss LSM 880 laser confocal system equipped with an Airyscan unit, environmental control chamber, and 1.4 NA Plan-Apochromat 63x DIC M27 oil objective. Similar to epifluorescence imaging procedures, cells were acclimated on the microscope stage for ∼30 min, while being illuminated by photosynthetic light. To limit any potential laser-induced stress on cells (and also changes to Vipp1 localization) we identified fields of view with the use 594 nm laser set to the lowest power at maximal detector gain. Image stacks were acquired for 16 Z-planes (512 x 512) with a voxel size 40 x 40 x 180 nm centered at mid-cell where the focus was the sharpest. Laser lines 458 nm, 515 nm and 594 nm were used through the same beamsplitter (MBS 458/514/594) for excitation of CurT-mTurquoise2, Vipp1-GFP (mGFPmut3 variant: excitation maximum - 501 nm, emission maximum - 511 nm) and thylakoid respectively. Emissions from the CurT and Vipp1 channels passed through a dual bandpass filter BP 420-480, BP 495-550). Pixel dwell time (set at the fastest) and laser power were adjusted to avoid saturation and bleaching effects while the detector gain was set to 800. For Airscan processing, Zen Black 2.1 was used to process the image stacks by performing the filtering, deconvolution and pixel reassignments in 3D mode at default settings. This processing confers the increased spatial resolution (by a factor of 1.5 to 1.7x both laterally and axially) and an enhanced signal to noise ratio (Korobchevskaya *et al.*, 2017).

For live-cell imaging, an aliquot of cells (obtained from an OD_750_ of 0.2 - 0.3 culture) was placed onto the glass of a 35 mm MatTek dish (P35G-0.170-14-C) (MatTek, Ashland, MA) under a thin BG11 - 1% agarose pad (2 mm thickness) prepared in advance and covered with a coverslip to reduce evaporation. The dish was placed onto the objective in the stage insert and around the agarose pad a basin of water was poured to create a humidified atmosphere. The temperature was maintained at 30 ° C while the photosynthetic light was provided by a white light LED ring: RL1360 or DF198 (Advanced Illumination, Rochester, VT) that was positioned on the stage insert and controlled by an analog signal through Zen 2.1. The LED white light was programmed to switch off only during the fluorescence image acquisition. The light intensity reaching the cells at the objective lens was 100 µE m^2^ s^-1^ which permitted a growth rate comparable to the rate measured in bulk culture. To limit photobleaching and phototoxicity, the imaging light power was limited to 20% by the FL attenuator and the light exposure for image acquisition in the GFP channel (20 to 30 ms) was adjusted to generate a signal-to-background ratio of less than 1.5. Also, the Z-series were limited to 12 steps or fewer. For thylakoid channel, all exposures used were 3.2 ms. Cell doubling time under these conditions was approximately 8 h.

Multispectral datasets were aligned using the affine transformation plugin in Zeiss Zen 2.1 using a standard prepared from TetraSpeck (Invitrogen, Carlsbad, CA) multicolor beads immobilized on a 1.5 coverslip.

To obtain filtered images, a custom MATLAB script was written to computationally enhance both diffraction-limited spots and contours of thylakoids by applying a local maxima filter that fit a 3D Gaussian to every position in the dataset and returned the Maximum Likelihood Estimate (MLE) of the amplitude, background and square-error. The amplitude component of the MLEs reports a quantitative visualization of the diffraction limited signal since it is separated from its background component.

To automatically segment cells in the brightfield channel a custom MATLAB script was written that implemented correlation imaging as described previously (Julou *et al.*, 2013). In summary, the edge of a cell is uniquely defined by its Z intensity profile given a 3D brightfield Z-stack. Therefore, to computationally find all cell edges, a brightfield Z-stack was filtered in Z by cross-correlating this unique Z-intensity profile using the MATLAB function *convn*. This cross-correlated dataset was then thresholded and all enclosed objects were then segmented by MATLAB’s function *regionprops*.

Fiji (Schindelin *et al.*, 2012) was used to prepare the kymograms, extract the intensity profiles, and convert the final images. Unless noted, in all images displayed, the grayscale contrast was auto-adjusted linearly before converting to bitmaps.

### Electron microscopy

*Synechocystis* cultures were grown to an OD_750_ of 0.2, incubated briefly in media supplemented with 100 mM mannitol and centrifuged. Aliquots of the pellet were high pressure frozen in a Wohlwend Compact 02 High Pressure Freezer at the University of Colorado-Boulder Electron Microscopy Facility and stored under liquid nitrogen. Frozen samples were then freeze-substituted at −80 C in 0.05% uranyl acetate in acetone and embedded in Lowicryl HM20 in a Leica AFS. For immunodetection, thin sections (50-60 nm) were labeled with rabbit polyclonal anti-GFP (generous gift of Pam Silver, Harvard University) diluted 1:500 in 1% non-fat dry milk in PBST, followed by secondary labeling with 5 nm gold-goat anti-rabbit Ig diluted 1:20 in PBST. Labeled grids were further contrasted with 2% aqueous uranyl acetate and lead citrate. Electron micrographs were obtained with an FEI Tecnai Spirit BioTwin TEM operating at 100 kV equipped with an AMT 2Kx2K side entry CCD camera.

To obtain transmission electron microscopy images of cells from dark- and light grown cultures presented in Fig. 6C and Fig. S6, the cells were collected by centrifugation, resuspended in a 100 µl of 20% BSA in BG-11 medium, loaded onto 100 µm-deep carriers and frozen in a Wohlwend Compact 01 High Pressure Freezer at the Electron Microscopy Facility at the HHMI/Janelia Research Campus. Samples were freeze-substituted in 2% osmium tetroxide/0.1 % uranyl acetate/3% water in acetone using a fast freeze-substitution method (McDONALD and Webb, 2011) and embedded in Eponate 12 resin. Ultrathin sections (60-65 nm) were post-stained with uranyl acetate/lead citrate and imaged in a Tecnai 12 electron microscope (FEI, Hillsboro, OR) operating at 80kV equipped with an Ultrascan 4000 digital camera (Gatan Inc, CA).

### Spectrophotometry

Whole-cell absorption and fluorescence emission spectra and were obtained on Spectramax i3 multimode reader (Molecular Devices, Sunnyvale, CA). For low-temperature measurements the cells were suspended in 50% glycerol and flash frozen in liquid nitrogen.

### Oxygen production measurements

Initial rates of oxygen production were obtained from dark- (144 hours) and light-growing (24 hours post-dark) cultures. Cells were diluted in BG11 medium supplemented with 10 mM NaHCO_3_ to an OD_750_ of 0.3, which corresponds to a cell density of ∼2.5 *10^8^ cells per ml, as estimated by a Multisizer 3 Coulter Counter (Beckman Coulter). Measurements were performed with an InLab^®^OptiOx sensor connected to the S9 Seven2Go Pro dissolved oxygen meter (Mettler Toledo) at 23° C, with moderate stirring in a closed vessel. Linear rates of oxygen production were obtained in complete darkness (respiration) or in light (photosynthesis) at 300 µE provided by a white light LED ring (DF198, Advanced Illumination, Rochester, VT).

## AKNOWLEDGEMENTS

We thank Courtney Ozzello and Thomas Giddings of University of Colorado-Boulder Electron Microscopy Facility for their immunoelectron microscopy work and discussions on the project. We also thank Amalia Passoli of HHMI/Janelia for her electron microscopy imaging of dark- and light grown cells and Eric Wait of Advanced Imaging Center at HHMI/Janelia for his help in generating the 3D rendering of Vipp1-GFP and CurT-mTurquoise2 expressing cells. We are grateful to Jeffrey Moffitt, Joseph Markson, and members of the O’Shea laboratory for their comments on the manuscript. This work was funded by the Howard Hughes Medical Institute. FC was supported by grants to Nancy Kleckner from the National Institutes of Health: RO1 GM044794 and RO1 GM025326.

---

**Fig. S1.**
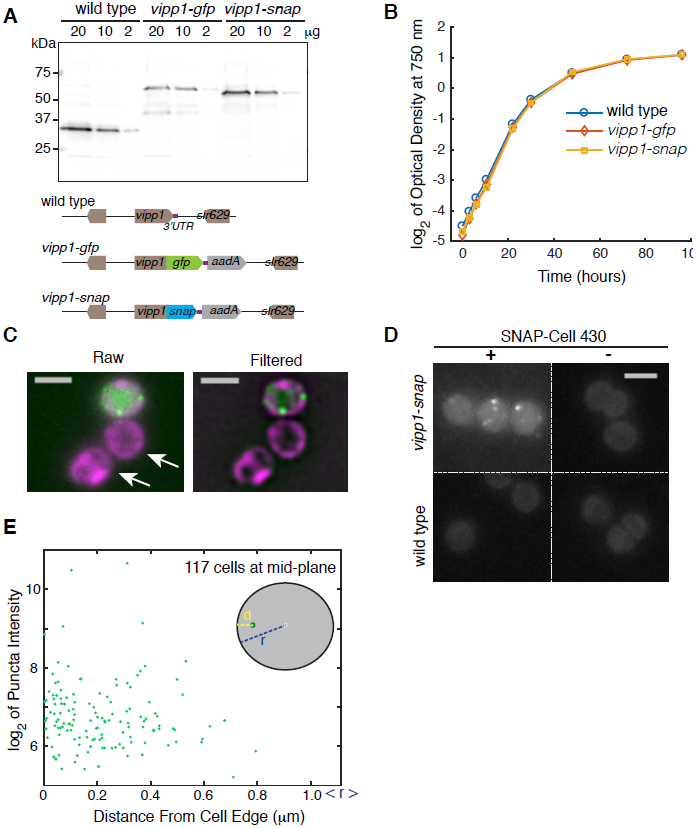
Characterization of Vipp1 expression and cellular localization in strains with epitope-tagged versions of Vipp1. **A**) Immunodetection of Vipp1 in cell lysates obtained from wild type, *vipp1-gfp* and *vipp1-snap* cells. Western blotting was performed as previously described (Gutu and O’Shea, 2013). Anti-Vipp1 rabbit polyclonal antibodies raised against Vipp1 from *Chlamydomonas reinhardtii* (Agrisera AS06145) were used. Below are the diagrams of the Vipp1-tagged DNA constructs used in this study (see Methods). **B**) Growth of wild type, *vipp1-gfp* and *vipp1-snap* strains in liquid culture bubbled with a mix of air and 1% CO_2_ at 100 µE intensity light. **C**) Merged Vipp1 (green) and thylakoid (magenta) images of Maximum Z-Intensity Projections of a mix of *vipp1-gfp* and wild type cells (arrows) showing the difference in cytosolic signal intensity (i.e. diffuse fraction of Vipp1) between the two cell types. On the right, an illustration of the filtering procedure (see Methods) used to enhance visualization of puncta and thylakoid features is shown. Scale Bar = 2 µm. **D**) Images of Maximum Z-Intensity Projections (raw epifluorescence) of wild type and *vipp1-snap* cells labeled with 5 µM SNAP-Cell 430 substrate (NEB S9109S) (after 20 min incubation in growth conditions) showing that Vipp1 puncta form independently of the GFP-tag. To facilitate comparison in signal intensity, the same grayscale contrast was applied to all four images. Scale bar = 2 µm. **E**) Intensities and distances of Vipp1 puncta relative to the cell edge. For any Vipp1 puncta found at or near the mid-Z plane of a cell, the intensity and the Euclidian distance (d) between its *xy* coordinates and the nearest point on the cell boundary edge obtained from segmentation in the brightfield channel was extracted and plotted. The <r> on the x-axis is the average cell radii of the 117 analyzed cells. Related to Fig. 1B for which the same dataset was used.

**Fig. S2.**
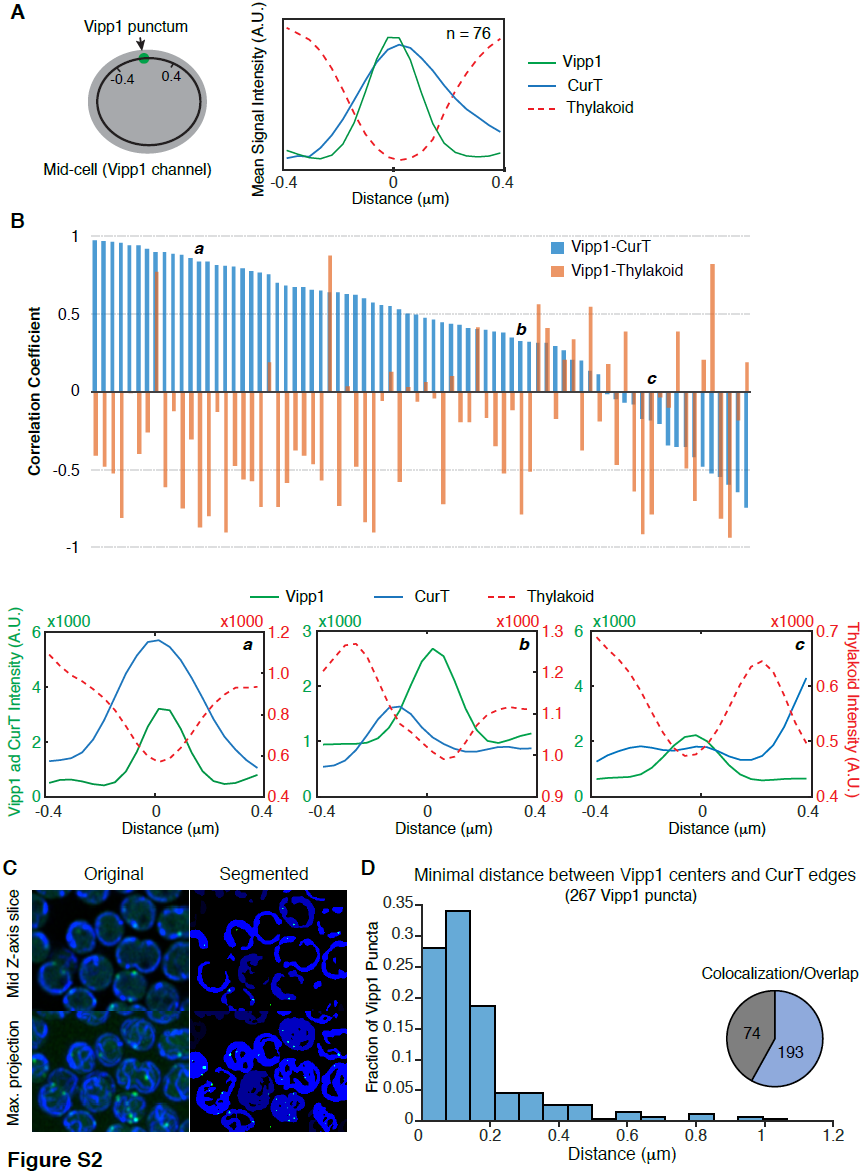
Vipp1 puncta colocalize with CurT enrichments at thylakoid edges. A) **Left** - diagram illustrating how the arc line intensity profiles of Vipp1, CurT and thylakoid signals were obtained. The grey oval represents the cell at mid-Z plane. The green dot is a Vipp1 punctum localized at the same mid-Z plane. A manually traced circumferential (black circle) line parallel to the cell boundary and traversing the green Vipp1 punctum was used to extract the profile intensity in the Vipp1, CurT and thylakoid channels. For any given Vipp1 punctum identified as a peak in the profile (using Matlab ’s *peakfinder* function), the intensities of the CurT and thylakoid signals were also collected for a 0.4 µm arc segment on either side of the Vipp1 peak (marked with a black notch). 76 arc segments (0.8 µm length) each centered on a given Vipp1 peak were extracted from 38 independent circumferential profiles that were obtained from 3D stacks of live-cells imaged in the Vipp1, CurT and thylakoid channels. (A single circumferential profile can produce one or more Vipp1 peaks). The cells were imaged in superresolution mode on a confocal system equipped with the Airyscan detector (Zeiss LSM 880) (Korobchevskaya *et al.*, 2017) (see Methods). **Right** – ensemble averaged arc segments of Vipp1, CurT and thylakoid signals. **B)** Bar graph of the Pearson ’s correlation between Vipp1 and CurT, and Vipp1 and the thylakoid signals for each of the 76 datasets. In general, Vipp1 and CurT signals correlate positively, whereas Vipp1 and thylakoid signals correlate negatively (aggregated *p*-values: 1.22 *10^-6^ and 5.57x10^-62^ respectively, null hypothesis tested being there is no relationship between the observed measurements). The weaker correlation values are obtained when the long axes of the thylakoid edges are quasi-parallel to the intensity profiles lines. Three correlation examples (labeled ***a***, ***b*** and ***c*** in the bar plot) of intensity profiles are shown in bellow. **C**) Object-based colocalization analysis of Vipp1 puncta with the CurT enrichments. Example of Airyscan merged images of Vipp1 (green) and CurT (blue) channels and the corresponding segmentation masks obtained by iterative thresholding of the 3D stacks as implemented in the 3D ImageJ Suite plugin (Ollion *et al.*, 2013). Both mid Z-slice and Maximum Intensity Projection along the Z axis are shown. **D**) Histogram of the distance distribution of each Vipp1 punctum to its most adjacent CurT object (Vipp1 puncta centers to CurT object edges). 267 Vipp1 puncta obtained from an image stack containing ∼70 cells were analysed. Note that this analysis includes all distances regardless if the Vipp1 centers are “inside “ or “outside “ of the CurT objects. The majority of the distances between Vipp1 puncta and CurT edges fall under the optical resolution resolution limit afforded by the Airyscan confocal imaging system (140 nm lateral and 400 nm axial). Inset pie chart shows how many of the Vipp1 puncta volumes overlap (in cyan) with the CurT objects.

**Fig. S3.**
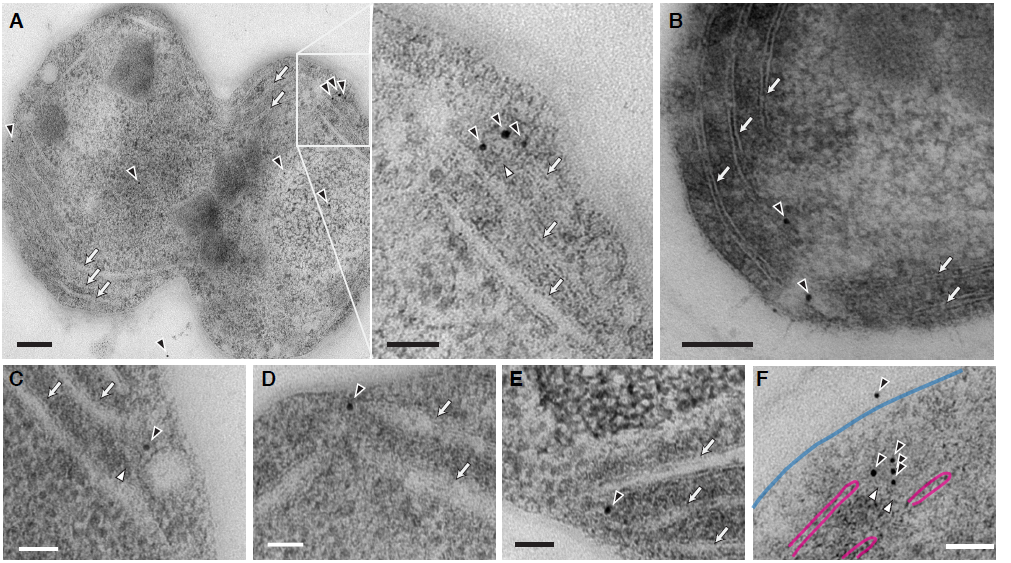
Detection of Vipp1-GFP by immuno-electron microscopy. Examples of Vipp1-GFP staining (black arrowheads) in cell sections obtained from *vipp1-gfp* strain. The thylakoid sheets (open arrows) run parallel to the plasma membrane forming several peripheral stacks per cell. The majority of the gold stain observed tend to localize at thylakoid edges which appear as tips in 2D images. Occasional ring-like structures (white arrowheads) of 30-35 nm in diameter could be observed at or near the gold particles at these regions (**A, C** and **F**). The ring-like structures at these sites are reminiscent of the previously described thylakoid centers (van de Meene *et al.*, 2006). Plasma and thylakoid membranes in panel **F** are indicated with cyan and magenta overlays. **A-B**: scale bar = 100 nm (**A** inset panel: scale bar = 50 nm). **C-F**: scale bar = 50 nm.

**Fig. S4.**
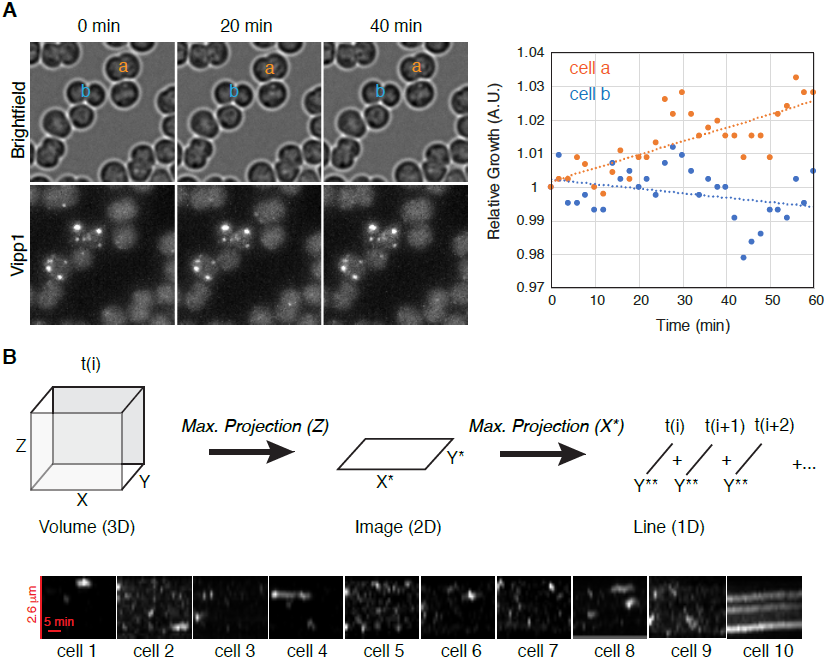
Punctuated dynamics of Vipp1 is a hallmark of all growing cells. **A**) **Left** - a montage of three time points from a 3D time lapse of growing cells, with brightfield on the top row and Vipp1 for the bottom row (Maximum Intensity Projections of epifluorescent 3D stacks - 8Z steps spaced every 280 nm). A small subset of the cells are non-growing (see cell labeled “b”) and in these cells Vipp1 exists predominantly as bright and long-lived puncta. Actively growing cells manifest diffuse Vipp1 signal and Vipp1 puncta that are dimmer and transient (see cell labeled “a”). **Right** - quantification of cell growth (as normalized cell areas relative to time 0) of cells “a” and “b” indicated in the montage on the left. Cell area measurements were extracted from automatic segmentation masks performed on the brightfield 3D stacks (see Methods). Cells were grown in light (100 µE m^2^ s^-1^) under an agarose pad, directly on the microscope stage and imaged every 2 min. Dotted trend lines represent least-square regressions of the raw measurements. **B**) To better visualize the formation and number of Vipp1 puncta over time of a given cell imaged in time-lapse mode, we made kymograms from 3D stacks of individual cells by two successive dimension reductions. First, a 2D image of Maximum Z-intensity Projection of a given cell was obtained, then the *x*-dimension was maximum projected onto the *y*-axis, which was eventually concatenated in a time series to make the kymogram. Here we show kymograms obtained from 10 representative cells in the Vipp1 channel (filtered) collected every 1 min for 30 min (8 Z-steps spaced at every 280 nm). Vipp1 puncta number and dynamics vary slightly from cell to cell - lasting generally 1-2 min. Occasionally (as also shown in panel A), we observed Vipp1 puncta failing to disassemble and slowly losing intensity due to photobleaching; they would occur in a non-growing cell (for example, cell 10).

**Fig. S5.**
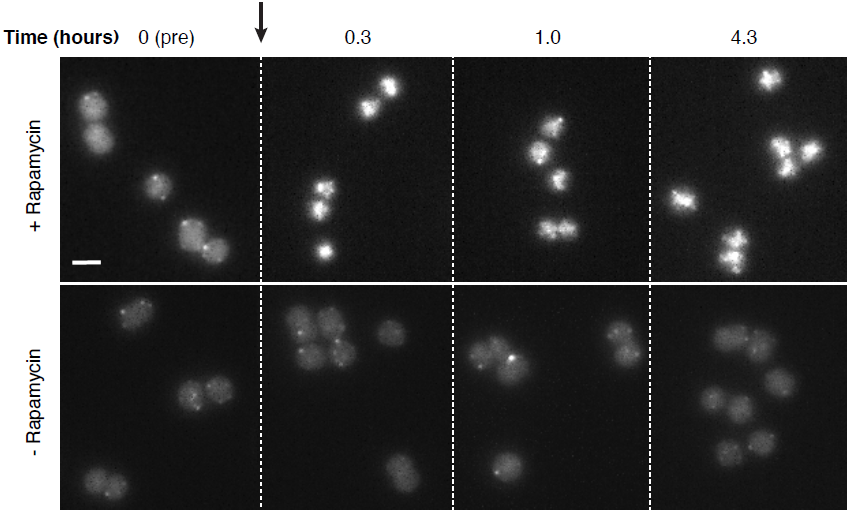
Time-course of rapamycin-induced Vipp1 relocalization. Maximum Z-Intensity Projection images of cells expressing all the components of the anchor-away system and taken from a log-phase culture bubbled with 1% air-CO_2_ mix at 100 µE light intensity at different times before and after addition of 20 µM rapamycin or solvent (dimethyl sulfoxide) alone showing that relocalization of Vipp1 to the nucleoid at the cell centers is rapid, effective and stable for at least 4-5 hours. The same grayscale contrast was applied to all panels within each condition.

**Fig. S6.**
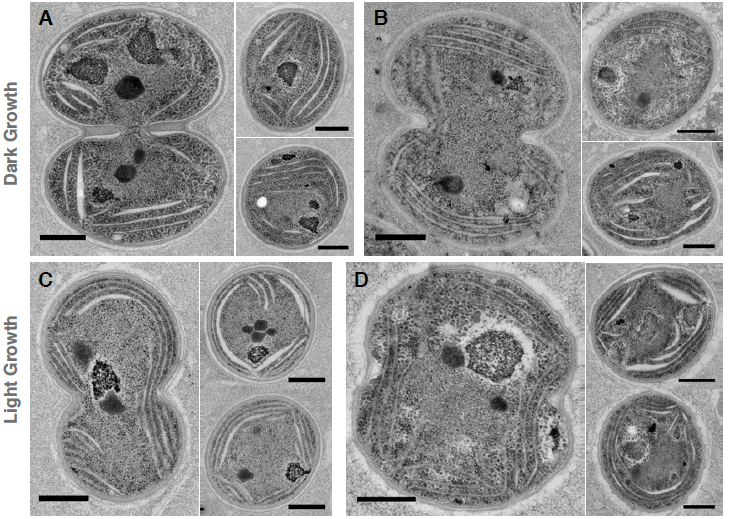
Representative transmission electron microscopy images of dark-and light-grown cells expressing all the components of the anchor-away system after 48 hours of growth in darkness or in light in the absence or presence of 20 µM rapamycin. Dark-grown cells obtained by culturing the cells for 144 hours in darkness in the presence of glucose with brief exposures to activating light (see Methods) were diluted to an OD_750_ of 0.1 in glucose-containing media and to which either dimethyl sulfoxide solvent (**A**) or 20 µM rapamycin dissolved in dimethyl sulfoxide was added (**B**) and further grown in complete darkness for 48 hours. Light-grown cells obtained from a log-growing culture were similarly diluted to an OD_750_ of 0.1 in media with no glucose to which dimethyl sulfoxide alone (**C**) or 20 µM rapamycin in dimethyl sulfoxide (**D**) was added and grown in light for 48 hours. The overall morphology of the thylakoid stacks in rapamycin-treated cells is similar to that of the control cells. Scale bars = 0.5 µm.

**Fig. S7.**
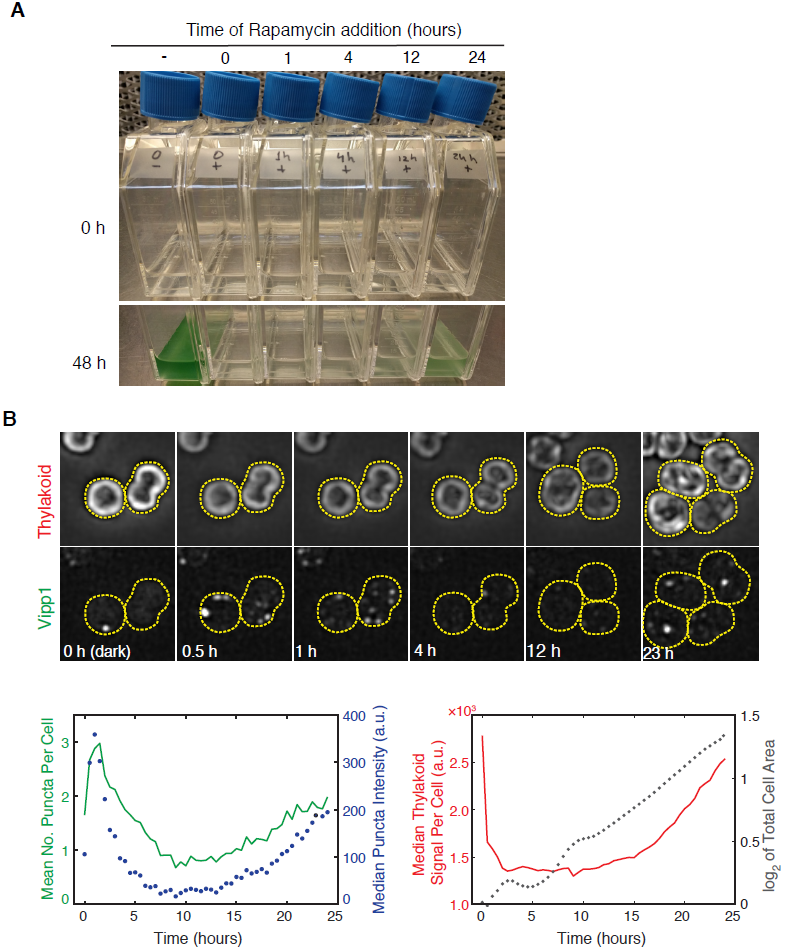
The early stages of *Dark-to-Light Transition* are more prone to the effect of Vipp1 perturbation. **A**) Delaying Vipp1 relocalization to a later time during the *Dark-to-Light Transition* lessens the impact of the perturbation on growth – as evidenced by the increased pigmentation (i.e. growth) of cultures labeled 12 and 24 h. **B**) Vipp1 undergoes a burst in the rate of puncta formation in the early stages of the *Dark-to-Light Transition*. **Top -** selected images from a time-lapse montage of dark-grown *vipp1-gfp* cells transitioning to *Light Growth*. Maximum Z-Intensity Projections of 3D filtered stacks in the thylakoid and Vipp1 channels are shown. The same grayscale contrast was applied to the entire montage within each channel. Bottom: quantification of Vipp1 puncta number per cell, Vipp1 puncta intensities, total thylakoid signal per cell and total cell area extracted from brightfield segmentation masks over time (every 30 min for 24 h) from a field ∼260 starting *vipp1-gfp* cells. At time 0 h, before light is turned on, thylakoid auto-fluorescence per cell is high, likely because the existing photosynthetic pigments residing in the thylakoid membranes are uncoupled from photochemical reactions (hence more fluorescent). In growing photosynthetic light, total thylakoid signal per cell is relatively flat early on and starts accumulating after ∼12 hours as the cells increase their thylakoid content and acclimate fully to the photosynthetic lifestyle. The initial dip in total cell area reflects the shrinkage of a small fraction of cells that fail to grow. For full movie of a representative group of cells - see Video S6.

**Video S1.** 3D rendering of a live cell expressing Vipp1-GFP and CurT-mTurquoise2 based on an image stack obtained by Airyscan imaging. Vipp1 signal was thresholded to isolate out the Vipp1 puncta (green), which colocalize with the CurT enrichments (blue) situated the edge of thylakoid sheets (red). The movie was created using a modified version of the 5D renderer used in (Wait *et al.*, 2014). Scale bar = 1 µm.

**Video S2.** Vipp1 puncta are short-lived and form in all growing cells. Time-lapse imaging (every 2 min for a total of 60 min) of *vipp1-gfp* cells growing photosynthetically directly on the microscope stage. Maximum Z intensity projections (8-Z slices spaced at 280 nm) of raw epifluorescent images in the Vipp1 channel are shown. Scale bar = 2 µm.

**Video S3.** Majority of Vipp1 puncta appear and disappear at the cell periphery on a timescale ∼1-2 min. Time-lapse imaging (every 30 sec for 10 min) of Vipp1 (left) and thylakoid (middle) channels of a group of photosynthetically growing cells. Merged filtered images of the middle Z-planes are shown. The arrow indicates an individual of Vipp1 punctum assembling and disassembling over time. Related to Fig. 3A.

**Video S4.** Vipp1 puncta have limited mobility. Continuous imaging in the Vipp1 channel (every 81 milliseconds for 8 sec) at mid-Z plane. The images were first bleach-corrected by histogram matching and then Laplacian-filtered to enhance visualization of Vipp1 puncta and cell edges. Related to Fig. 3C.

**Video S5.** Vipp1 relocalization during the *Dark-to-Light Transition* induces a severe growth defect. Timelapse movie (every 20 min for 24 h) of a mix of *vipp1-gfp-fkbp12* cells - with or without the HU-FRB anchor - transitioning from dark to light and to which 20 mM rapamycin was added after the first timepoint to induce Vipp1 relocalization. Maximum Z-Intensity Projections (10 Z-steps) of raw epifluorescent images in the thylakoid (left), Vipp1 (middle) and brightfield (right) channels are shown. The photosynthetic light (100 µE m^2^ s^-^1) was turned on after the first timepoint. The cells in the which the Vipp1 signal coalesced at cell center (i.e. cells expressing the HU-FRB anchor) fail to grow or grow slower than the cells in which Vipp1 remains diffuse with occasional peripheral puncta (no HU-FRB anchor). Related to Fig. 5C.

**Video S6.** The early stages of the *Dark-to-Light Transition* are marked by a burst of Vipp1 puncta formation. Time-lapse movie of *vipp1-gfp* cells transitioning from *Dark* to *Light* growth. Dark-grown cells were placed under an agarose pad prepared with glucose-free media and imaged every 30 min for 24 h in brightfield, thylakoid and Vipp1 channels. Photosynthetic growth light was turned on after the first timepoint. Filtered Maximum Z-Intensity Projections (9 Z-steps) are shown for the thylakoid (left) and Vipp1 (right) channel. The linear grayscale contrast remained unchanged for the entire time series within each channel. Related to Fig. S7B.

